# Improving classification of correct and incorrect protein-protein docking models by augmenting the training set

**DOI:** 10.1101/2022.10.22.512683

**Authors:** Didier Barradas-Bautista, Ali Almajed, Romina Oliva, Luigi Cavallo, Panos Kalnis

## Abstract

Protein-protein interactions drive many relevant biological events, such as infection, replication, and recognition. To control or engineer such events, we need to access the molecular details of the interaction provided by experimental 3D structures. However, such experiments take time and are expensive; moreover, the current technology cannot keep up with the high discovery rate of new interactions. Computational modeling, like protein-protein docking, can help to fill this gap by generating docking poses. Protein-protein docking generally consists of two parts, sampling and scoring. The sampling is an exhaustive search of the tridimensional space. The caveat of the sampling produces a large number of incorrect poses, producing a highly unbalanced dataset. This limits the utility of the data to train machine learning classifiers. Using weak supervision, we developed a data augmentation method that we named hAIkal. Using hAIkal, we increased the labeled training data to train several algorithms. We trained and obtained different classifiers; the best classifier has 81% accuracy and 0.51 MCC on the test set, surpassing the state-of-the-art scoring functions.

## 1. Introduction

Protein-protein interactions (PPIs) are fundamental to life, as they are at the basis of many biological processes. Correct and specific interactions between proteins are required for proper functioning within organisms, and it is now accepted that perturbation of these interactions can lead to defective phenotypes (Sahni *et al*., 2015; Cheng *et al*., 2021). This explains why the development of new drugs also rests on manipulating PPIs (Lu *et al*., 2020). This background highlights the importance of knowing the structure of protein-protein complexes at atomic level. Unfortunately, the number of experimental structures of protein complexes corresponds to a small fraction of the characterized PPIs (Mosca *et al*., 2013). A solution to this problem consists in the prediction of the structure of protein-protein complexes using molecular docking approaches (Lensink *et al*., 2018; Harmalkar and Gray, 2021).

A docking process starts with the generation of 10^3^-10^5^ alternative three-dimensional (3D) decoys consisting in different interaction between the proteins in the complex, and is followed by a scoring step aimed at singling out the correct solutions within the ensemble of generated decoys. Proper scoring is thus a key step, object of blind assessment in a separate challenge of the CAPRI (Critical Assessment of PRedicted Interactions) experiment (Lensink *et al*., 2007).

Different approaches are used to develop scoring functions. The most popular are either energy-based (Moal *et al*., 2015; Cheng *et al*., 2007; Chaudhury *et al*., 2010) or knowledge-based (Huang, 2014; Moal *et al*., 2013; Vangone *et al*., 2017). Recently, hybrid scoring functions combining the above approaches (Vreven et al. 2011) (Pierce and Weng, 2007; Andrusier *et al*., 2007), or adding to them evolutionary information (Andreani *et al*., 2013) have been proposed. Finally, scoring functions based on the consensus of contacts between residues at the interface of the complex have been also developed (Chermak *et al*., 2016; Oliva *et al*., 2013). Despite all these efforts, the available scoring functions still have challenges in sorting out correct decoys from the generated decoys ensemble (Lensink *et al*., 2007).

This explains the need for developing novel and better scoring algorithms, potentially combining in a proper way the single scoring functions currently available (Barradas-Bautista *et al*., 2017), also by taking advantage of machine learning (ML) tools (Barradas-Bautista *et al*., 2022; Cao and Shen, 2020; Geng *et al*., 2020; Moal *et al*., 2017; Wang *et al*., 2020). One of the greatest challenges in developing novel scoring approaches consists in the limited number of correct decoys in the datasets used for training and testing. In fact, correct solutions in the decoy ensembles provided by docking often represent a tiny fraction of them. In many cases there are only a few dozen or even less correct decoys in an ensemble of thousands incorrect decoys. Coupled with the small number of available experimental protein-protein complex structures, this allows building datasets for training and testing novel scoring functions with only a relatively small number of entries. This clearly limits the possibility of applying advanced ML tools. To solve this challenge, we decided to increase the size of the datasets used to develop scoring functions using data augmentation techniques.

These techniques present an affordable, fast and available alternative to produce more training data. Data augmentation is the process of extracting additional useful information from existing data (van Dyk and Meng, 2001). It is very useful in expanding available datasets in terms of size and quality of labels. Such techniques are crucial, especially when generating new data is not viable due to cost, time, need of experts and access to special materials and facilities. Therefore, different fields are adopting relevant augmentation approaches. In the imaging field, image rotation, cropping, translation and other forms of pixel manipulations are applied to diversify and increase the size of the training dataset, which results in better classification accuracy (Wang *et al*., 2017). In other works, more complex augmentation techniques are employed, such as the use of conditional Generative Adversarial Networks (GANs) (Qian *et al*., 2019), to generalize the dataset into a broader feature domain. In addition, data augmentation has further proven its usefulness in more complex data representations, such as time series speech recognition data (Park *et al*., 2019) and medical imagery.

Currently the available curated data on the protein pair interactions including 3D structures is very limited, spanning only couple of hundreds of examples, while the number of reported binary PPIs ranges for several organism the hundreds of thousands (Szklarczyk *et al*., 2019). Only for the former set of PPIs, it is possible to assign a label of correctness to generated docking decoys, based on their measured similarity to the real 3D structure. This poses a challenge for the field to produce useful and inexpensive methods to augment a relevant training dataset. Fortunately, there are several works that suggest novel ways to focus on generic and automated methods of training data generation leveraging different techniques such as weak-supervision (Cubuk *et al*., 2019). Weak supervision techniques are being adopted as a data augmentation option to remediate labeled data scarcity. Using weak supervision to automatically generate data labels based on certain domain knowledge and heuristics is becoming common.

Snorkel is a weak-supervision framework (Ratner *et al*., 2020), successfully used on text mining and natural language processing (NLP) tasks. The framework applies statistical modeling on user-defined labeling functions (LFs) to infer class labels. LFs are simple methods that read-in data input and produce a confident-guessed class label or abstain. Then, it models and estimates the correlations and accuracy of each defined labeling function, producing an enhanced collective prediction model used to label unlabeled data.

In a recent work by Wang et al. (Wang *et al*., 2019), weak supervision was used to automatically label clinical text data, saving human effort. Furthermore, in a work by Fries et al. (Fries *et al*., 2019), they rely on domain experts to define heuristics to noisily label cardiac MRIs, which is consequently used as training data enhancing the detection of aortic valve malformations. The defined heuristics are integrated as labeling functions where the Snorkel statistical modeling is applied to generate class labels. This solution is used to automatically label the abundant sets of unlabeled MRIs, which are then used to train and enhance aortic valve malfunction classifiers. An example of the application of Snorkel in biology is to improve the retrieval of information from the literature and clustering families of genes according to ontology terms (Dutta *et al*., 2020). In biomedicine/biochemistry, Snorkel was used for extracting chemical reaction relationships from biomedical literature abstracts (Mallory *et al*., 2020). Both examples use Snorkel to widen the training of their respective machine learning approaches but are constraint to the use of textual heuristics.

In this work, we leverage docking decoys scoring functions information as the basis for setting up the Snorkel framework (Fig. 2). In our approach we explore the use of noisy encoding using only discrete values. The docking decoys can be theoretically classified using the results of scoring functions given that there is a correlation between the quality of the decoy and the underlying assumptions of the scoring function. We adapt each scoring function to be encoded as a Snorkel labeling functions, giving only numerical thresholds We provide further details on how we automated and defined the threshold in the section “Optimizing Snorkel”.

**Figure 1.**
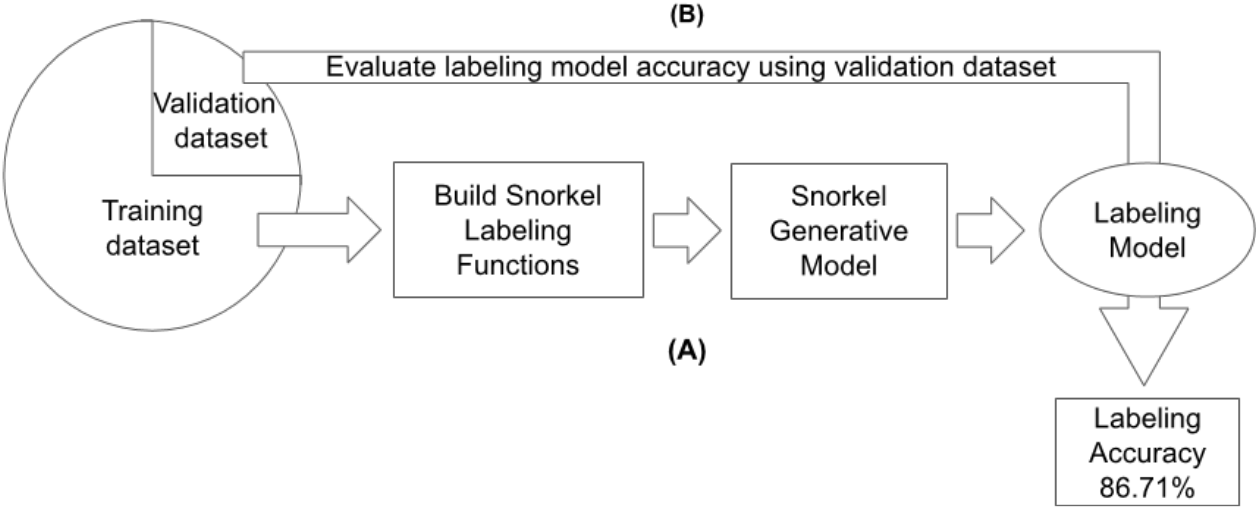
Building and fine-tuning the snorkel labeling model. We optimize Snorkel with our custom defined labeling functions, which are optimized with thresholds extracted from the training data set(A). Then the framework output quality is measured by its accuracy in labeling the validation dataset (B).

**Figure 2.**
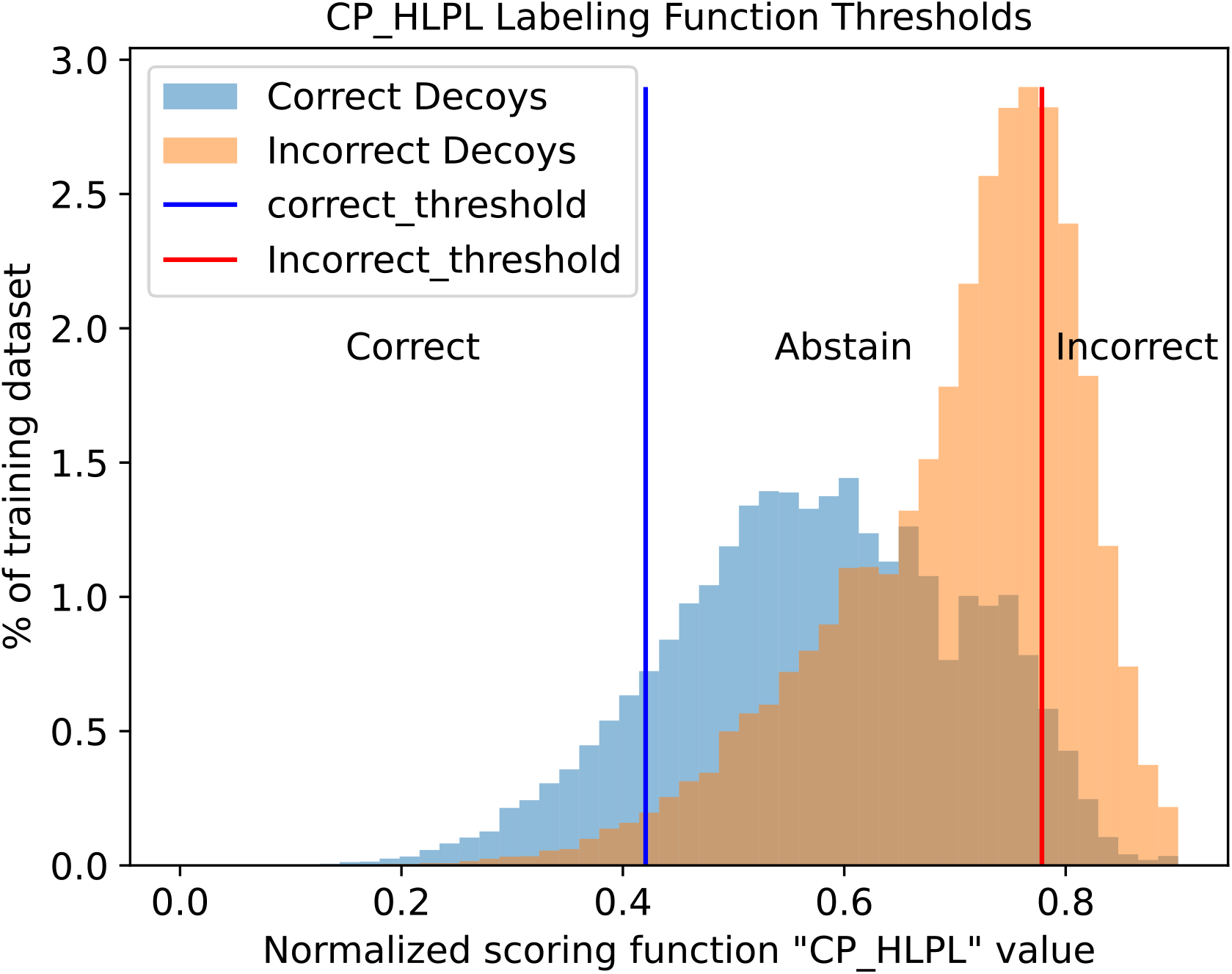
Histogram of training set, classified by CP_HLPL scoring function, showing the selected threshold for correct (blue line), incorrect (red line) and abstain (in between) used in the corresponding snorkel labeling function

## 2. Materials and Methods

### 2.1 Machine learning models and evaluation metrics

We performed data analysis and employed ML algorithms within the scikit learn python library (Pedregosa *et al*., 2011; Varoquaux *et al*., 2015; Buitinck *et al*., 2013) and within pyplot/seaborn for visualization (Hunter, 2007; Waskom *et al*., 2014). We obtained a single-layer Perceptron (PRC), a deep neural network model under Tensorflow 2 framework (TF), and a random forest (RF) classifier. We deem appropriate these three classifiers given that combinations among different scoring features could involve complicated associations since they can work at different levels of abstractions (coarse or atomic level) and under different physicochemical definitions where contact energy can consist of several terms with different weights.

PRC is a one-layer neural network; RF consists in a set of 100 decision trees, TF has a total of five layers, three layers to process the data, and two dropout layers, to minimize the possibility of overfitting. Further details on the classifiers implementation can be found in supplementary section 4.

We calculated global standard metrics to evaluate the ML predictions in our validation and test sets as follows:

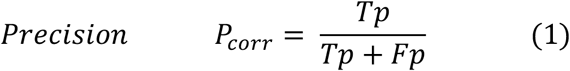

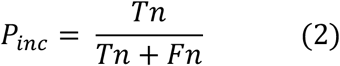

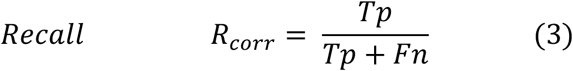

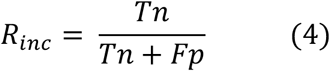

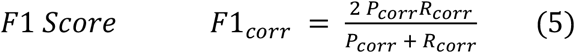

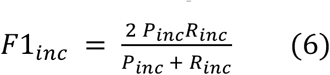

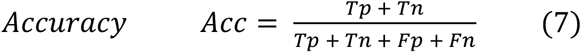

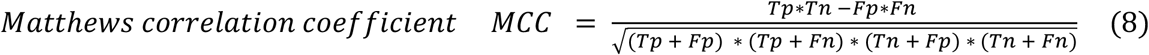

where Tp stands for true positives, Tn for true negatives, Fp for false positives, Fn for false negatives. For the independent testing on the Score_set (see below), we also used the success rate, that is the number or percentage of correct solutions within the top N positions.

### 2.2 Scoring functions (features)

We used a total of 157 descriptors (features) from public sources, analogously to what we did previously (Barradas-Bautista *et al*., 2022). Among the used features, 92 come from the CCharPPI server (Moal *et al*., 2015) and mostly consist in physics-based or empirical energy terms; 32 are calculated by our tools CONSRANK (Oliva *et al*., 2013, 20, 2015) and COCOMAPS (Vangone *et al*., 2012, 2011), and consist of the consensus CONSRANK score and of the number of interresidue contacts per class of involved amino acids; 28 come from CIPS (combined interface propensity per decoy scoring (Nadalin and Carbone, 2018), and represent sums and averages of the CIPS score over the different classes of residues at the interface; 3 represent the buried surface area (BSA) calculated with FreeSASA (Mitternacht, 2016), and 2 are non-interacting surface (NIS) terms (Kastritis *et al*., 2014), polar and apolar, calculated by Prodigy (Vangone and Bonvin, 2017). For details see Barradas-Bautista *et al*., 2022 (Supplementary data, Table S2). The above features were used to train the ML algorithms. The values of theses scoring functions were normalized as detailed in Barradas-Bautista *et al*., 2022. For each of the scoring functions, we performed a minmax normalization [0 - 1] on the training set, and applied the produced normalization on the test, validation and the silver sets.

Additionally, from each of the three trained classifiers with the original/”core” data set, Bal-BM4 set (see below), we obtained the “class probability” (probability to be correct) associated to each DM. We included these values as the last three features for the data set; since they values already range between 0 to 1, we did not apply any normalization here.

### 2.3 Docking models and datasets

#### 2.3.1 Generation and quality assessment of docking models (DMs) in the “core” sets

For “core” set hereon we mean the ensemble of docking decoys whose quality (label of correctness) could be trustworthily assigned based on the comparison with the real 3D structure of the corresponding targets. The starting point for the generation of our “core” set was the protein docking Benchmark 5 (BM5) (Vreven *et al*., 2015) consisting of 230 protein-protein complexes (targets) complete with their experimental structure. For each of the 230 protein-protein complexes (targets) in BM5 (Vreven *et al*., 2015), we generated a total of 30,000 DMs with FTDock (Gabb *et al*., 1997), ZDock (Chen and Weng, 2002) and HADDOCK (de Vries *et al*., 2007; Dominguez *et al*., 2003) as detailed previously (Barradas-Bautista *et al*., 2022). The quality of the generated DMs was assessed following the CAPRI (Critical Assessment of PRedicted Interactions) protocol (Méndez *et al*., 2003), and consequently DMs were classified, in order of increasing quality, as Incorrect, Acceptable, Medium- and High-quality, as reported in Table S1. All the DMs and relative quality assessment are available at (https://doi.org/10.5281/zenodo.4012018). The 17 targets for which no correct DM was identified were removed; for the 213 remaining targets, we classified overall 246 high quality, 7,146 medium quality, 42,815 acceptable and 6,339,793 incorrect DMs. For each target, we discarded DMs with a number of clashes greater than the average number of clashes plus two standard deviations for that target. Overall we discarded 243,061 DMs, which left us with a total of 6,146,939 DMs, on average 28,859 ±218 DMs per target. For further details can be found in pour previous work (Barradas-Bautista *et al*., 2022).

#### 2.3.2 Balanced dataset for ML analysis

Since databases containing DMs generated by docking software are highly unbalanced, to train the ML algorithms we built a “balanced” dataset by applying a random undersampling selection of the incorrect DMs to match the number of correct ones. The balanced dataset, Bal-BM5, features for each target the same number of correct and incorrect DMs. This number varies variable from 2 to 600 (on average 346 ±358) for the different targets, as it depends on the number of correct DMs available for each specific target. For details see previous work (Barradas-Bautista *et al*., 2022). The targets and DMs selected in the above steps were separated into training and validation sets using the following strategy. All the 161 targets included in the benchmark 4 (BM4) were assigned to the training set, Bal-BM4, while the remaining 52 targets, corresponding to the BM5 update, relatively to BM4, were assigned to the validation set, Bal-BM5up.

#### 2.3.3 3K unbalanced dataset for ML analysis

For the unbalanced datasets, we selected 3,000 DMs per target, out of those generated in the docking step and that passed the check on clashes. For each target we selected up to 600 DMs with acceptable or better quality, and picked up randomly incorrect DMs until 3,000 DMs in total were collected into the 3K-BM5 dataset. We fixed a maximum value for the number of correct DMs, 600, to avoid biasing the training of the ML algorithms towards a few cases with a too large number of correct DMs. A strategy similar to that used to split the whole Bal-BM5 dataset into training and validation datasets was used here to extract a validation dataset from the 3K-BM5. Specifically, the 52 targets corresponding to the BM5 update were assigned to the 3KBM5up validation dataset.

#### 2.3.4 Generation and quality assessment of docking models (DMs) in the “augmented” training set

In order to augment our training dataset, we started from the 1392 protein pairs from the high confidence human interactome in the I3D database (Mosca *et al*., 2013), to create a “silver set” that we then added to the above balanced core set to obtain our “augmented set”.

We used FTDock and ZDock to generate DMs for each of the 1392 protein pairs, with the settings reported above. A 3K decoys set per target was obtained as followed. The top 1000 DMs per target were selected from each program, according to their in-build functions. Moreover, following a strategy also used by Mosca for high-throughput docking experiments (Mosca *et al*., 2009), we selected additional 1000 DMs per target, by re-ranking the ZDock DMs with PyDock, and maintaining the top DMs according to this other scoring function.

On average, there are 201.3 ±87.1 overlapping DMs per target in the selection from the ZDock and PyDock scoring functions. In total, this represents 20.1 % of overlap between the two sets.

The quality label to each of these decoys was assigned by the Snorkel labeling model described below. After the evaluation, this silver set consisted of 4,168,957 DMs, 998.8 ±0.41 DMs for FTDock, 995.6 ±0.50 DMs for ZDock, and 996.6 ±0.50 for the ZDock/PyDock, on average per target. In the silver set, 1,767,386 DMs were labeled as correct (42.4%) and 2,401,570 DMs (57.6%) were labeled as incorrect.. On average, each of the 1392 protein pairs in the silver set has 1726.5 ±692.5 DMs labeled as correct and 1289.1 ±681.0 DMs labeled as incorrect.

We then combined the silver set with the above balanced core training dataset (161 targets included in BM4), to obtain our augmented training set. The augmented set thus consists of a total of 1553 protein pairs, with a total of 4,224,740 labeled DMs, being 76 times bigger than the original training set; each protein pair has on average 1171.8 ±731.7 DMs labeled as correct and 1565.4 ±810.8 DMs labeled as incorrect. All tabular data is available at https://repository.kaust.edu.sa/handle/10754/666961.

#### 2.3.5 Optimizing Snorkel

To find useful scoring functions that we could incorporate as LFs, we handle this problem as a binary classification problem by categorizing CAPRI quality standards “A”, “M”, “H”, as correct, and “I” as incorrect. Treating this problem as a binary classification allows us to establish a classical confusions matrix with the definition of True Negative (Tn), True Positive (Tp), False Positive (Fp), False Negative (Fn).

#### 2.3.6 Thresholds for labelling functions

After evaluating all the selected DMs with over 150 scoring functions and corresponding labels, we obtained the receiver operating curve (ROC) for all the possible scoring functions. Initially, we tried to train the Snorkel labeling model using all the available labeling functions, however according to the built-in snorkel analysis we observed many labeling functions overlapped on the coverage of the data producing conflicts ending on label accuracy equal to a random. Therefore, we decided to select a subset of labeling functions that perform best individually. We prioritize the selection based on the area under the curve (AUC) from the ROC analysis of each function. For each scoring function, we produced the ROC of all significant thresholds and calculated the AUC. Then we sorted all scoring functions by their AUC, which provides us an indication of how well they evaluate DMs individually. This analysis resulted in the selection of 40 different scoring functions that we could potentially use as labelling functions.

#### 2.3.7 Defining the labelling functions

Optimally we look for scoring functions that could provide a clear distinction between correct and incorrect DMs. Unfortunately, this is part of the problem we are attempting to solve. Thus, we target the best of what each scoring function could provide. Each scoring function evaluating a docking decoy produces a numerical result. We look for a certain threshold where the accuracy of this scoring function is very high in evaluating either correct and incorrect DMs to employ it as a labeling function.

The LFs that we aim to produce need to provide the best possible answer on a subset of the data, but not necessarily all of it, as abstaining is an option. Therefore, for the correct DMs, we aim to maximize R_corr_ (eq. 3) and minimize the false-positive recall, R_Fp_, (eq. 9).

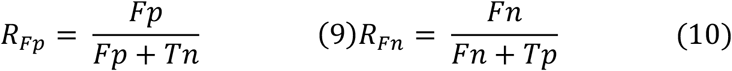

Also, for the incorrect DMs, we aim to maximize R_inc_ and minimize the false-negative recall, R_Fn_ (eq.10)). Therefore, we calculate two ratios: the “CorrectThreshold” as R_corr_ over R_Fp_, eq. 11, and the “Incorrect Threshold” as the R_inc_ over R_Fn_, eq. 12.

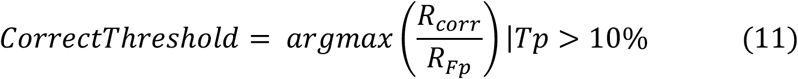

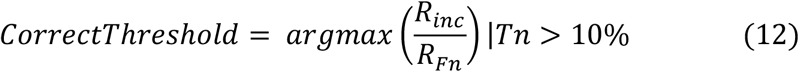

If the value of the scoring function does not fall within either threshold, it is considered as an abstain. The determination of the threshold was a useful metric to ascertain the utility of the scoring function as LFs. This analysis is similar to feature selection with a wrapper-like approach Kohavi and John (1997) (Kohavi and John, 1997); John *et al*. (1994) (John *et al*., 1994) (Fig. S6). We selected only those LFs with a coverage equal or greater than 20% total consisting of at least 10% correct, and at least 10% incorrect samples. (Fig. 3 and Table S3). By setting the coverage on 10% for both labels on each LF produced a total coverage on 68% on the training dataset. The final selection consisted in eight labeling functions that produced the best accuracy for labelling the training set (Table S3) at the cost of reducing the coverage but avoiding a high overlap between the LFs.

**Figure 3.**
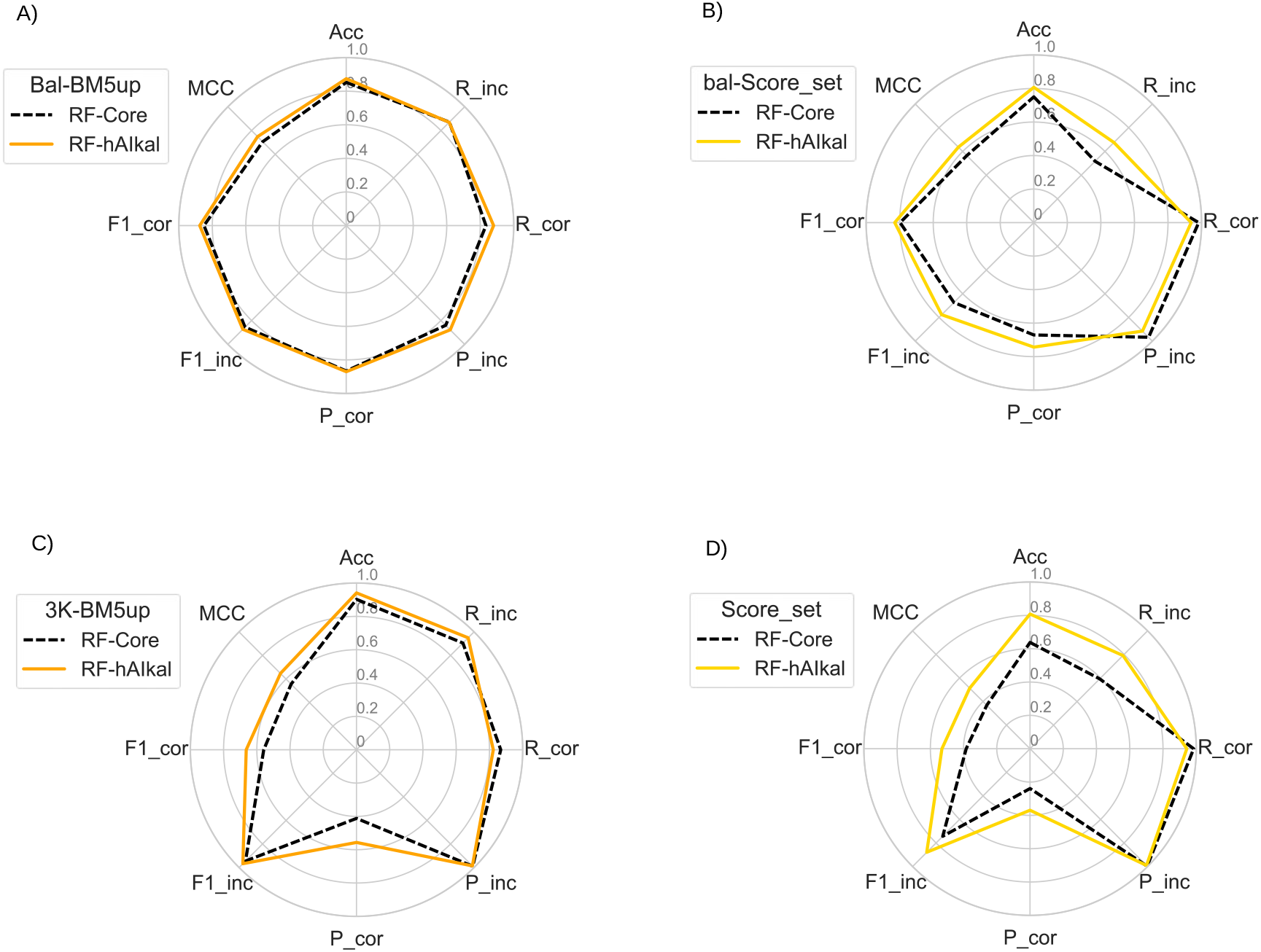
Radar plots of the performance metrics for the RF classifiers when trained with the “core” (RF-core) or augmented (RF-hAIkal) data on **A-B**) the balanced validation (Bal-BM5up) and test (Bal-Score_set) sets, and **C-D**) the unbalanced validation (3K-BM5up) and test (Score_set) sets.

#### 2.3.8 Feature selection and weak supervising model labeling

Using a Pearson correlation analysis, we can see that many of the scoring functions have a positive correlation (Fig. S7). Some of the scoring functions have previously shown synergy combined with many other scoring functions, such as CP_HLPL and CP_TSC (Barradas-Bautista *et al*., 2017). Meanwhile, there are also anti correlations, such as the case of BSA_apolar. This finding might be related to the biochemistry of the interfaces. Many of the scoring functions are built upon electronic potentials, whereas apolar patches are an essential part of the interface composition (Levy, 2010). Also, others are collinear, or derivations such as CIPS_AlAr and AlAr, which are respectively a pair potential combining interface composition with Aliphatic-Aromatic residue contact preference and the total count of aliphatic and aromatic residue contacts. CONSRANK_val is not strongly correlated with most of the scoring functions, despite being the best scoring function with the highest AUC. This benefits the ML models since this scoring function is orthogonal to the others.

The Snorkel framework allows us to inspect the performance on the training set for possible conflicts between labeling functions and obtain the labeling’s accuracy. We trained the Snorkel labeling model with our data, and we ensure the maximum coverage possible with the least conflict possible. We tested the produced weak-supervision labeling model, and we observe that it achieves 86.3% labeling accuracy on the training dataset and 86.7% labeling accuracy on the validation dataset (Bal-BM5up).

We applied the resulting labeling model to the DMs in silver set (see above).

### 2.4 CAPRI Score_set as an independent test set

For an independent testing, we used the CAPRI Score_set, made of 15 targets from the CAPRI Rounds 13-26, with a total of 19,013 DMs, predicted by 47 different predictor groups including web servers, which use different docking and scoring procedures, being thus highly diverse (Lensink *et al*., 2016). DMs are complete of quality label and only 2,166 (11%) are classified correct (acceptable or better). This is therefore our “unbalanced test set”. Analogously to what done above with BM5, we also derived from Score_set a balanced dataset, our “balanced test set”, where only the 13 targets with at least one correct solution were included and a random selection of the incorrect DMs was applied to match the number of correct ones for each target. This resulted in a set of 3990 DMs, equally distributed between correct and incorrect ones.

## 3. Results

### 3.1 Design of the classifiers

We chose to train the classifiers on a balanced dataset because this would allow us to better learn the class represented by the DMs in the data set. Herein we applied a data augmentation strategy to design a machine learning classifier of predicted 3D structures of PPIs (docking models, DMs, in the following). To this aim, we added to a “core” set of 55748 DMs for 161 PPIs in BM4 (Vreven *et al*., 2015), for which a correctness label was available based on their similarity to the real 3D structure, a “silver” set of over 4 × 10^6^ for 1392 PPIs from the I3D database (Mosca *et al*., 2013), to which the quality label was assigned by a Snorkel label model here developed (described in detail in the Methods section).

Three machine learning algorithms, single-layer Perceptron (PRC), a deep neural network model under Tensorflow 2 framework (TF), and a random forest (RF) one, were used to train a classifier for correct/incorrect DMs. Classifiers were trained with a balanced version (same number of correct and incorrect DMs) of either the “core” or the augmented set (respectively 55748 and 4224704 DMs). Optimal values for the parameters obtained for each ML algorithm by training with the “core” data were maintained in the “augmented data” classifiers. For these classifiers, the “class probability” (probability to be correct) associated with the DMs by the three “core” classifiers were used as additional features (see Methods). Hereon, classifiers trained with “core” data will be named PRC/TF/RF–core, while those trained with the augmented data will be named PRC/TF/RF–hAIkal.

In the following, the feature selection for the classifiers is discussed; then results of the testing of the obtained classifiers on different datasets, balanced and unbalanced, with the latter being more representative of a “real case scenario”, are presented and discussed. The comparison in performance for the classifiers obtained with or without augmented data is emphasized. In addition, the classifiers performance is compared to that of some classical scoring functions for DMs available in literature (Moal *et al*., 2015). For the performance evaluation, global standard metrics were used: the Matthews’ correlation coefficient (MCC), accuracy (Acc) and, calculated both for the correct and incorrect DMs, recall (R_corr_, R_inc_), precision (P_corr_, P_inc_) and F1_score (F1_corr_, F1_inc_).

### 3.2 Feature selection for training models at different stages

We selected 40 scoring functions (the top 25%) that we could potentially use as labeling functions in the Snorkel model, based on an AUC threshold of 0.67 calculated on BM4-bal. These scoring functions include knowledge-based potentials at an atomic or residue level (such as AP_DARS, AP_MPS, CP_HLPL, CP_SKOIP, CP_TB, CP_TSC), energy-based functions (e.g. PyDock and SIPPER)(Cheng *et al*., 2007; Pons *et al*., 2011), and other descriptors of the interaction (e.g. buried surface area, number of the aliphatic and aromatic contacts), as well as the class probabilities from each of our machine learning models, featuring as expected the highest performance, with an average AUC value of 0.97 (±0.02). The entire list with description is given in Table S2.

Of the 157 descriptors from public sources, the one having by far the highest AUC value, 0.92, was the CONSRANK score. This is a consensus score, which reflects the conservation (or frequency) of the inter-residue contacts featured by a given model in the whole decoys ensemble it belongs to (Chermak *et al*., 2015; Oliva *et al*., 2015, 2013), widely tested on CAPRI targets (Barradas-Bautista *et al*., 2020; Lensink *et al*., 2019, 2016; Vangone *et al*., 2013). The second highest AUC value, 0.78, was obtained for AP_GOAP_DF, an atomic distance-dependent potential, the DFIRE term in the GOAP energy (Zhou and Skolnick, 2011), that we obtained from the CCharPPI server (Moal *et al*., 2015). Next, four scoring functions, CP_HLPL, CP_MJ3h, CP_RMFCA, and DDG_V, shared the same AUC value, 0.77; the first 3 being residue contact/distance-dependent potentials, the fourth one being a miscellaneous scoring function, consisting in a microscopic surface energy model derived from mutation data (Moal *et al*., 2015; Feng *et al*., 2010; Rajgaria *et al*., 2006).

For the final optimization of the Snorkel model, we selected eight LFs with minimum overlap. They included the the six scoring functions with the top three AUC values, CONSRANK score, AP_GOAP_DF, CP_HLPL, CP_MJ3h, CP_RMFCA, DDG_V, plus two LFs from machine learning models developed in this work (from NNC and TF). As we selected the RF model for training the final classifier, we didn’t include the RF class probability within the selected LFs. The final labeling model obtained from these eight LFs improved the labeling accuracy to 86% on the Bal-BM4 set and to 88% on the Bal-BM5 set.

### 3.3 Baseline classifiers performance on the Bal-BM5up validation dataset

We used Bal-BM5up (BM version 5 update) as a validation dataset. This is a balanced dataset featuring only PPIs not included in the BM4 we used for the training. This was done to avoid a bias in the performance evaluation due to the fact that some of the features selected for the classifiers training were actually developed using parts of the BM4.

On this dataset, all classifiers have better than random metrics. The MCC values are indeed ≥ 0.70, while all the other metrics considered range between 0,83 and 0.91 (Tables 1 and S5). Overall, the best performance is exhibited by the RF classifier trained with augmented data, which features the maximum values for most of the metrics. Use of augmented data improves instead only the recall for correct and precision for incorrect DMs for the perceptron and the TF classifiers, as compared to the same classifiers trained with “core” data.

**Table 1.** Classifiers performance on Bal-BM5up (balanced validation set). “Core” and hAIkal correspond to the algorithms trained with the “core” and augmented data, respectively. The highest number for each metric is highlighted in bold.

|  | PRC-core | PRC-hAlkal | TF-core | TF- hAlkal | RF-core | RF- hAlkal |
| --- | --- | --- | --- | --- | --- | --- |
| MCC | 0.7104 | 0.6954 | 0.7311 | 0.7228 | 0.7059 | **0.7489** |
| Acc | 0.8552 | 0.8446 | 0.8655 | 0.8612 | 0.8527 | **0.8744** |
| F1<sub>corr</sub> | 0.8564 | 0.8543 | 0.8663 | 0.8636 | 0.8499 | **0.8751** |
| F1<sub>inc</sub> | 0.8539 | 0.8334 | 0.8647 | 0.8586 | 0.8554 | **0.8738** |
| R<sub>corr</sub> | 0.8635 | **0.9117** | 0.8717 | 0.8791 | 0.8343 | 0.8797 |
| R<sub>inc</sub> | 0.8468 | 0.7774 | 0.8594 | 0.8432 | **0.8711** | 0.8692 |
| P<sub>corr</sub> | 0.8493 | 0.8038 | 0.8611 | 0.8487 | 0.8662 | **0.8706** |
| P<sub>inc</sub> | 0.8612 | **0.8980** | 0.8701 | 0.8746 | 0.8402 | 0.8784 |

The three algorithms have similar performance in terms of accuracy, which could indicate either the upper limit of the prediction capacity on this balanced testing set or that some part of the data is noise. Moreover, given that a binary classification is a strict correct and incorrect division, we might misclassify borderline DMs in a “grey zone” where some of our features give conflicting scores.

### 3.4 Classifiers performance on the independent balanced test set, Bal-Score_set

Our Bal-Score_set, as detailed in the Methods section, consists of 3,990 DMs, equally distributed between correct and incorrect, that we derived as a subset of the public Score_set benchmark for 15 targets from the CAPRI Rounds 13-26 (Lensink *et al*., 2016). Such DMs were generated by 47 different predictor groups including web servers, with different docking and scoring procedures, thus featuring a wide diversity. On this test set, which is independent of the targets and DMs we used in the training process, the evaluation metrics are quite variable but still outline an overall better than random performance for all of them (Tables 2 and S6). While the MCC values vary in a limited range, from 0.50 to 0.64, *indicating a similar overall performance*, the other metrics range between close-to-random (see the R_inc_ for the PRC-hAIkal and RF-core classifiers) to close-to-perfect prediction values (see R_corr_ and P_inc_ for all the classifiers). Again, RF-hAIkal has overall the best performance with the highest values for all the eighth metrics, but the R_corr_ and P_inc_. For the latter two metrics, the highest values were still achieved by a hAIkal classifier, specifically PRC-hAIkal.

**Table 2.** Classifiers performance on Bal-Score_set (the balanced test set). “Core” and hAIkal correspond to the algorithms trained with the “core” and augmented data, respectively. The highest value for each metric is highlighted in bold.

|  | PRC-core | PRC-hAikal | TF-core | TF- hAikal | RF-core | RF-hAikal |
| --- | --- | --- | --- | --- | --- | --- |
| MCC | 0.6082 | 0.4947 | 0.6248 | 0.5992 | 0.5652 | **0.6363** |
| Acc | 0.7784 | 0.7018 | 0.7937 | 0.7723 | 0.7502 | **0.8070** |
| F1<sub>corr</sub> | 0.8155 | 0.7687 | 0.8239 | 0.8117 | 0.7975 | **0.8295** |
| F1<sub>inc</sub> | 0.7227 | 0.5804 | 0.7511 | 0.7121 | 0.6741 | **0.7777** |
| R<sub>corr</sub> | 0.9794 | **0.9910** | 0.9643 | 0.9809 | 0.9829 | 0.9382 |
| R<sub>inc</sub> | 0.5774 | 0.4125 | 0.6229 | 0.5634 | 0.5171 | **0.6757** |
| P<sub>corr</sub> | 0.6986 | 0.6278 | 0.7192 | 0.6923 | 0.6709 | **0.7434** |
| P<sub>inc</sub> | 0.9656 | **0.9786** | 0.9457 | 0.9672 | 0.9680 | 0.9160 |

**Table 3.** Classifiers performance on the 3K-BM5up dataset (unbalanced validation set). “Core” and hAIkal correspond to the algorithms trained with the “core” and augmented data, respectively. The highest value for each metric is highlighted in bold.

|  | PRC-core | PRC-hAikal | TF-core | TF- hAikal | RF-core | RF-hAikal |
| --- | --- | --- | --- | --- | --- | --- |
| MCC | 0.5523 | 0.4717 | 0.5790 | 0.5401 | 0.5565 | **0.6473** |
| Acc | 0.9088 | 0.8569 | 0.9064 | 0.8887 | 0.9024 | **0.9407** |
| F1<sub>corr</sub> | 0.5626 | 0.4647 | 0.5771 | 0.5343 | 0.5585 | **0.6639** |
| F1<sub>inc</sub> | 0.9491 | 0.9174 | 0.9474 | 0.9368 | 0.9451 | **0.9675** |
| R<sub>corr</sub> | 0.8239 | 0.8732 | **0.8974** | 0.8968 | 0.8677 | 0.8228 |
| R<sub>inc</sub> | 0.9154 | 0.8556 | 0.9071 | 0.8881 | 0.9050 | **0.9498** |
| P<sub>corr</sub> | 0.4271 | 0.3166 | 0.4253 | 0.3805 | 0.4118 | **0.5564** |
| P<sub>inc</sub> | 0.9855 | 0.9888 | **0.9914** | 0.9912 | 0.9889 | 0.9859 |

A comparison of the performances on this dataset of the random forest classifiers trained on the “core” and the augmented data is shown as radar plots in Figure 3.

### 3.5 Classifiers performance on the unbalanced validation and test sets, 3K-BM5up and Score_set

3K-BM5up and Score_set, a validation and a test set, respectively, are both highly unbalanced, thus allowing a *bonafide* assessment of the classifiers performance in close to real-world scenarios. Singling out the correct DMs from these hardly unbalanced datasets is particularly challenging. In particular, 3K-BM5up includes a total of 157,767 DMs, of which 92.8% are incorrect and only 7.2% are correct, while Score_set has 19,011 DMs, of which 88.5% incorrect and 11.5% correct.

Starting from 3K-BM5up, all the metrics indicate a better than random performance, with the exception of the P_corr_ for most classifiers (then reflecting into the F1_corr_ values). The precision in retrieving correct DMs (P_corr_) is clearly affected by the small number of relevant items (correct DMs) in the dataset. However, the recall in retrieving them (R_corr_) features quite satisfying values, above 0.82 for all the classifiers. This implies that, while including some incorrect DMs in the correct class, the classifiers can still recognize a large fraction of the relevant items, which is, in fact, the most desirable outcome.

MCC values explore a limited range, between 0.47 and 0.65, here RF-hAIkal is in general the best metrics for the prediction of the correct DMs. Although, R_corr_ is not the highest of all the classifiers, the F1_corr_ and MCC values indicate that RF-hAIkal has the better tradeoff of recall-precision and quality of prediction for the correct DMs. At first glance, on the 3K-BM5up set (unbalanced validation), the perceptron_hAIkal has the lowest values among the three classifiers. As expectd for unbalanced sets, all the three algorithms successfully classify many decoys according to accuracy. The accuracy of 86%, 89%, and 94% for perceptron_hAIkal, TF-hAIkal, and RF-hAIkal (Supplementary Table S7), respectively, is higher than the accuracy observed on the Bal-BM5up (balanced validation). The high accuracy can be associated to the strong influence of the larger number of incorrect decoys. Thus, the F1-score can show which algorithms have the best retrieval rate for correct decoys. Using this metric, RF-hAIkal has the best F1-score with 0.66.

While the balanced test set, Bal-Score_set, already evidenced a diversity in the classifiers’ performance, its unbalanced version, Score_set, further emphasizes it. MCC ranges between 0.29 and 0.51, staying above a random performance. As for the other metrics, the accuracy ranges between 0.52 for the PRC-hAIkal to 0.80 for the RF-hAIkal. P_corr_ is quite low (below 0.37) for all the classifiers, and consequently, the F1_corr_ explores a range around random values (between 0.32 and 0.53). However, similarly to the case of the 3K-BM5up, recall values for correct DMs are very high (above 0.94 for all the classifiers). RF-hAIkal has overall the best performance on this dataset as well, featuring the highest values for six metrics and staying very close to the maxima (and above 0.94) for the remaining two, i.e. R_corr_ and P_inc_. Most relevant improvements over the other classifiers concern recall for incorrect decoys, R_inc_, (0.79, to be compared to the 0.59 of the same algorithm trained on the core set) and, consequently, F1_inc_ (0.81 vs. 0.64 of RF-core). Although intrinsically low, P_corr_ is also significantly higher for this classifier.

A comparison of the performance on this dataset of the random forest classifiers trained on the core and the augmented data is shown as a radar plot in Figure 3B.

### 3.4 Effect of data augmentation on the classifiers performance

The main scope of this work is exploring the possible beneficial effect of data augmentation for the training of classifiers of PPIs DMs. Therefore, in the following, we will compare the performance of classifiers based on the same algorithms but trained respectively with the “core” or augmented data we set up here. Since, as we will see, the effect of data augmentation was particularly relevant to the RF classifiers, performance metrics for the RF-core and RF-hAIkal classifiers on the different datasets are shown and compared in Figure 3.

#### 3.4.1 Balanced datasets

Starting from the baseline performance on the balanced validation set (Bal-BM5up), we observe that the performance of the TF and PRC classifiers was virtually unaffected by the data augmentation, with variations in the MCC and accuracy values within 0.015. We notice however that data augmentation significantly improved the recall for correct solutions for the PRC-hAIkal classifier, which features indeed the maximum value overall. Looking at the RF-hAIkal classifier, instead, we observe a relevant improvement of the performance over RF-core, reflected in an increase of the MCC value from 0.71 to 0.75. With the exceptions of R_inc_ and P_corr_, staying mostly unaffected, the RF-hAIkal improves in fact all the metrics by 0.02-0.04.

The effect of data augmentation becomes more apparent, while preserving a similar trend, when the classifiers are tested on the balanced test set (Bal-Score_set). While the performance of TF was only marginally decreased (MCC from 0.63 to 0.60), that of PRC was more significantly affected, with the MCC changing from 0.61 to 0.50. However, the R_corr_ and P_inc_ metrics were improved again, by ≈0.02, for both the neural network-based algorithms. As for the RF classifiers, due to the use of augmented data during training, the MCC raised from 0.57 to 0.64 and accuracy increased from 0.75 to 0.81. All the metrics were indeed improved for the RF-hAIkal classifer (with increases up to 0.1) but for the R_corr_ and P_inc_, which were actually decreased while staying pretty high in value.

#### 3.4.2 Unbalanced datasets

With its 157,767 DMs, 3K-BM5up is the largest among the datasets we used for testing (≈9-fold larger than its balanced version and 8.4 times bigger than the Score_set). The effect of data augmentation here was similar to that previously observed on the Bal-Score_set. The performance of the TF classifier was only marginally affected (MCC decreased from 0.58 to 0.54 and accuracy from 0.91 to 0.89), while that of the PRC classifier was significantly decreased (MCC from 0.55 to 0.47 and accuracy from 0.91 to 0.86). However, R_corr_ was again significantly increased, from 0.82 to 0.87, for the PRC-hAIkal classifier.

The unbalanced independent test set (Score_set) is clearly the most challenging dataset for the classifiers testing. It indeed emphasizes the effect of data augmentation while substantially preserving the already observed trends. On this set, both the neural network-based algorithms decreased their performance, although, again, it was the PRC classifier to be more significantly affected (with MCC decreasing from 0.41 to 0.29 and accuracy from 0.69 to 0.52). On the other hand, the RF-hAIkal demonstrated a substantially improved performance. The RF-hAIkal MCC value increased to 0.51 *vs* the 0.27 of RF-core and the accuracy was 0.81 *vs* the 0.64 of RF-core. Values of all the metrics were significantly increased (by 0.13 to 0.2), but for R_corr_ and P_inc_, which remained virtually unaffected. Once more, the RF algorithm performance was boosted with the augmented data set.

### 3.5 Classifiers performance on the unbalanced sets in terms of success rate and comparison with other scoring functions

The success rate is the number (or fraction) of targets for which at least a correct solution can be predicted at the top N positions of a ranking and is a gold standard metric for the performance of scoring functions and algorithms for PPI DMs. We calculated the success rate over the top-1, top-10 and top-100 ranked positions for the classifiers we developed and for the 157 scoring functions we initially used as features (see Methods). The unbalanced validation and test sets, 3K-BM5up and Score_set were selected for such testing and comparison. Data for the six classifiers is reported in Table 5, while a comparison for the 10 top performing methods overall, in terms of top-10, is shown in Figure 4 (values for the top-20 performing methods are given in Table S5 and S6).

**Table 4.** Classifiers performance on the unbalanced Score_set (unbalanced test set). “Core” and hAIkal correspond to the algorithms trained with the “core” and augmented data, respectively. The highest value for each metric is highlighted in bold.

|  | PRC-core | PRC-hAikal | TF-core | TF- hAikal | RF-core | RF-hAikal |
| --- | --- | --- | --- | --- | --- | --- |
| MCC | 0.4106 | 0.2903 | 0.4916 | 0.4224 | 0.3685 | **0.5137** |
| Acc | 0.6930 | 0.5179 | 0.7802 | 0.7047 | 0.6376 | **0.8080** |
| F1<sub>corr</sub> | 0.4214 | 0.3191 | 0.5030 | 0.4333 | 0.3845 | **0.5305** |
| F1<sub>inc</sub> | 0.7911 | 0.6268 | 0.8589 | 0.8003 | 0.7432 | **0.8793** |
| R<sub>corr</sub> | 0.9811 | **0.9917** | 0.9671 | 0.9815 | 0.9843 | 0.9431 |
| R<sub>inc</sub> | 0.6560 | 0.4570 | 0.7559 | 0.6687 | 0.5925 | **0.7905** |
| P<sub>corr</sub> | 0.2683 | 0.1902 | 0.3399 | 0.2780 | 0.2389 | **0.3691** |
| P<sub>inc</sub> | 0.9963 | **0.9977** | 0.9944 | 0.9964 | 0.9966 | 0.9907 |

**Table 5.**
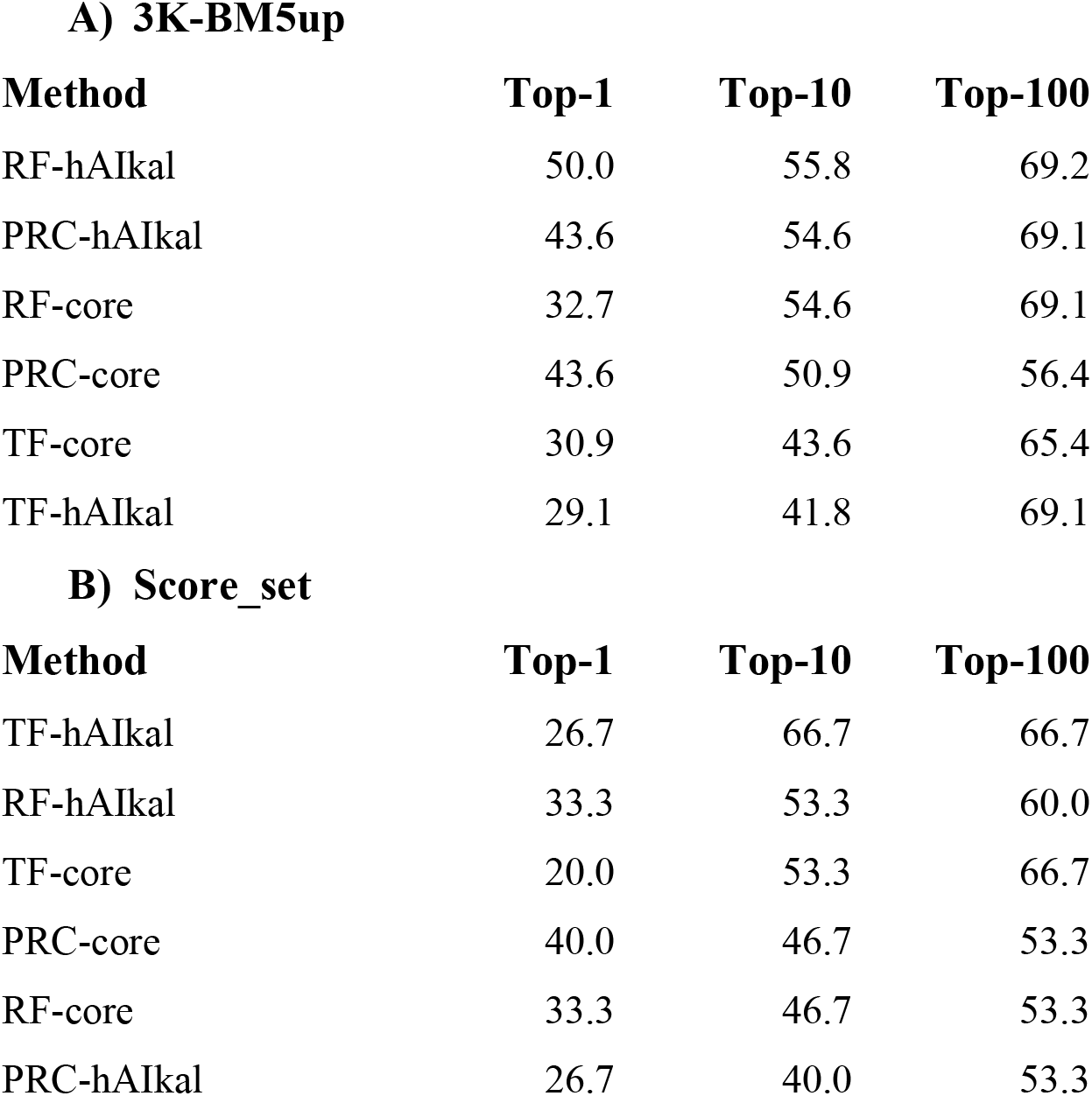
Success rate for the six here developed classifiers in terms of percentage of targets featuring at least one correct solution within the top-1, top-10 and top-100 ranked positions, sorted by top-10.

| **A) 3K-BM5up** |  |  |  |
| --- | --- | --- | --- |
| **Method** | **Top-1** | **Top-10** | **Top-100** |
| RF-hAlkal | 50.0 | 55.8 | 69.2 |
| PRC-hAlkal | 43.6 | 54.6 | 69.1 |
| RF-core | 32.7 | 54.6 | 69.1 |
| PRC-core | 43.6 | 50.9 | 56.4 |
| TF-core | 30.9 | 43.6 | 65.4 |
| TF-hAlkal | 29.1 | 41.8 | 69.1 |
| **B) Score_set** |  |  |  |
| **Method** | **Top-1** | **Top-10** | **Top-100** |
| TF-hAlkal | 26.7 | 66.7 | 66.7 |
| RF-hAlkal | 33.3 | 53.3 | 60.0 |
| TF-core | 20.0 | 53.3 | 66.7 |
| PRC-core | 40.0 | 46.7 | 53.3 |
| RF-core | 33.3 | 46.7 | 53.3 |
| PRC-hAlkal | 26.7 | 40.0 | 53.3 |

**Figure 4.**
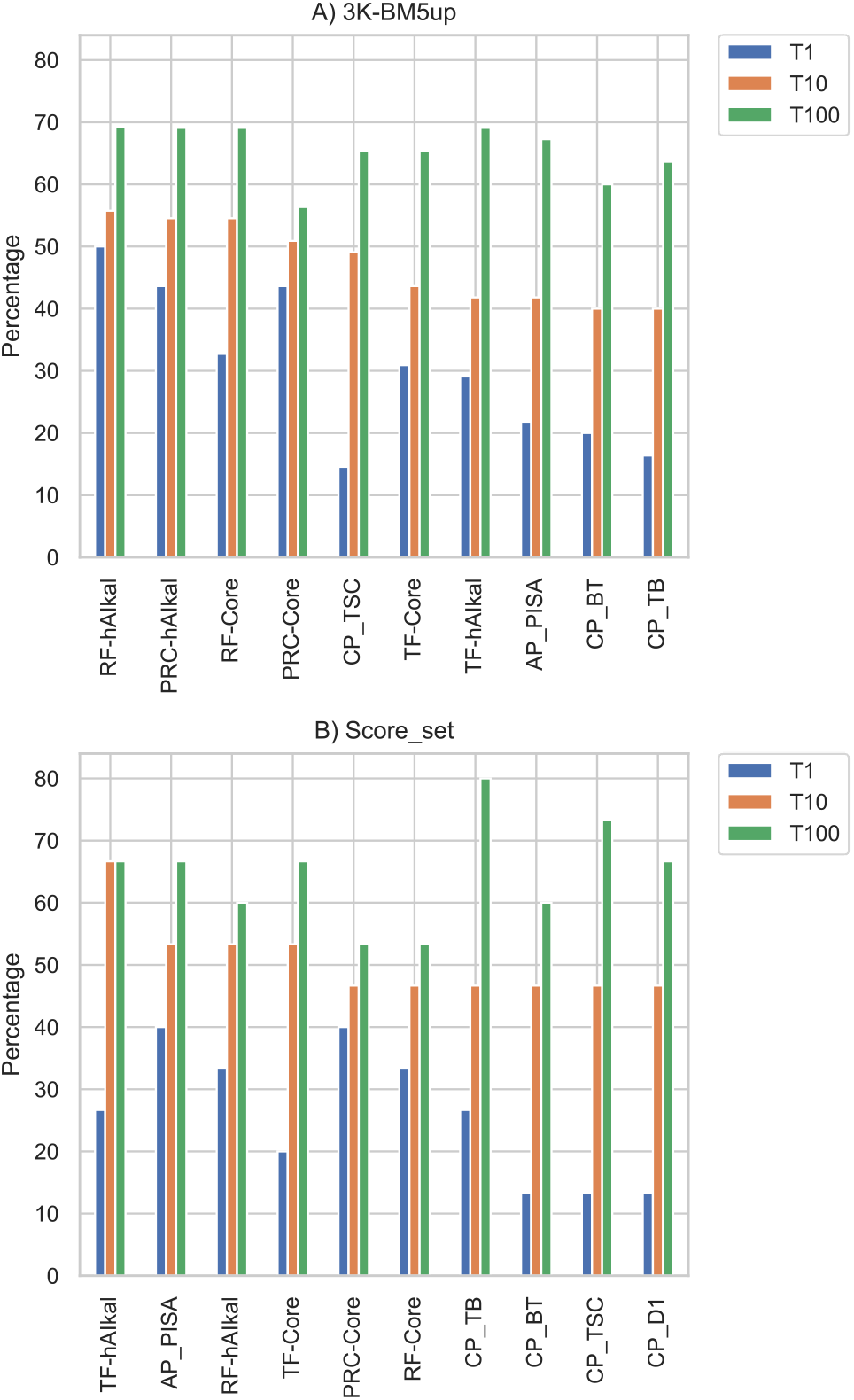
Percentage of BM5-up (**A**) and Score_set (**B**) targets for which at least 1 correct decoy was ranked within the top-1, top-10 and top-100 positions. Results are shown for the 10 best performing predictors among the here developed classifiers and considered scoring functions (see Methods), sorted by top-10.

On the 3K-BM5up (Table 5A), RF-hAIkal and PRC-haikal have the highest success rate in all rankings. The top-10 ranking of the classifiers trained with the “core” data is close to the hAIkal-trained classifiers. This trend seems to indicate that hAIKal data allows the classifiers to have a better performance. On Score_set, among the here developed classifiers (Table 5B), TF-hAIkal is the top performing one, with two thirds of the targets (67%) featuring at least one correct solution at the top-10 ranked positions. TF-hAIkal achieves indeed the top performance overall among the methods we tested.

In the overall comparison of the classfiers and scoring functions on the 3K-BM5up, TF-hAIkal is then followed by RF-hAIkal, TF-core and AP_PISA, with 53% targets and by the three “core” classifiers and four residue contact potentials, with a percentage of 47%. PRC-hAIkal occupies the 11^th^ position of this ranking, with a top-10 of 40% and a top-1 of 27%, likewise the scoring function CONSRANK (Table S5).

Therefore, the six classifiers we developed are all among the top performing methods on this independent testing, together with an atomic potential (AP_PISA, (Viswanath *et al*., 2013)) and four residue contact potentials (CP_TB, CP_BT, CP_TSC, CP_D1, (Feng *et al*., 2010; Tobi, 2010; Liu and Vakser, 2011; Tobi and Bahar, 2006).

Furthermore, we also obtained the success rate for the Score_set (unbalanced test set) (Figure 4B). On top-1 success rate, the PRC-core is tied with AP_PISA (Viswanath *et al*., 2013) (40%), RF-hAIkal and RF-Ori have the second-best success rate for top-1 ranking (33%), TF-hAIkal and PRC-haikal are on the third-place ties with two other scoring functions CONSRANK_val and CP_TB (27%). With a broader range of rankings, the differences between the various scoring functions are smaller The top-100 ranking demonstrates that it is worth exploring different approaches, such as non-deterministic scoring functions that show expanded use for particular cases of PPIs (Table S6). In this ranking, TF-hAIkal and TF-core, with a success rate of 67%, outperform RF-hAIkal (60%). However, both are tied with state-of-the-art scoring functions such as AP_PISA.

## 4. Conclusions

In this work we used CCharPPI and in-house scoring functions to train popular machine learning algorithms combined with weakly supervised learning. Overall, the trained machine learning models present high accuracy and high success rate over test and validation sets, surpassing reported methods or stand-alone scoring functions. Compared to other methods reported, we show that our approach increases the performance of the chosen algorithm. Unlike other methods based on CNNs with limited data to only one method (Balci *et al*., 2019; Wang *et al*., 2020) our approach expands the amount of data available for training by integrating up to three well-known PPI methods.

We developed a different way to use continue values to produce categorical data using a weakly supervised learning. The data augmentation using Snorkel and the threshold optimization quickly allowed the labelling of thousands of data points. The augmentation method worked best for a RF algorithm rather than for neural network ones. Meanwhile, the RF algorithm resulted to be more robust to noise from the data, and also less prone to outfitting, than neural network ones, coming on top as the best classifier. The difference between the neural networks can be explained from the architecture point of view. It is possible to change this outcome by using different architectures for neural networks either altering the drop-off layers or increasing the number of neurons or layers. Furthermore, we provide an extensive machine learning training data set, by using weak-supervised labeling of more PPIs. This allows covering a larger fraction of the PPIs space, and the developed data set could be used for developing novel AI-based strategies.

## Notes

https://repository.kaust.edu.sa/handle/10754/666961

https://doi.org/10.5281/zenodo.4012018

### Competing Interest Statement

The authors have declared no competing interest.

