## supplemental_material for "Improving classification of correct and incorrect protein-protein docking models by augmenting the training set"

**Table S1** . The Capri quality definitions, depend on different combinations of LRMSD, IRMSD and fnat. For most of the categories there is a need to fulfill one of the two conditions indicated on each columns.

| Quality | Condition 1 | Condition 2 |
|---|---|---|
| Incorrect | LRMSD > 10 Åand IRMSD > 4 Å fnat < 0.1 | fnat < 0.1 |
| Acceptable | (0.1 < fnat < 0.3) and (LRMSD ≤ 10 Å or IRMSD ≤ 4 Å) | fnat ≤ 0.3 and (LRMSD > 5 Å or IRMSD > 2 Å) |
| Medium | 0.3 ≤ fnat < 0.5) and (LRMSD ≤ 5.0 Å or IRMSD ≤ 2 Å | fnat ≥ 0.5 and (LRMSD > 1.0 Å and IRMSD > 1.0 Å) |
| High | fnat ≥ 0.5 and (LRMSD ≤ 1 Å or IRMSD ≤ 1 Å) | |

**Table S2** List of selected features used for the analysis and training. For each feature, the public source, relative reference and a short description are provided.

| Feature | Source | Description | Reference |
|---|---|---|---|
| **CONSRANK_val** | CONSRANK | CONSRANK normalized score, reflecting the conservation of the inter- residue contacts of a given model in the decoys set | Bioinformatics 31:1481 (2015) |
| **AP_GOAP_DF** | CCharPPI | The DFIRE term in the GOAP energy | Biophys J. 2011 101(8): 2043-2052. |
| **CP_MJ3h** | CCharPPI | Contact potential, calculated between intermolecular residues | Proteins 59(1):49 (2005) and BMC Bioinformatics 11:92 (2010) |
| **DDG_V** | CCharPPI | A microscopic surface energy model derived from mutation data | Proteins 83(4):640 (2015). |
| **CP_HLPL** | CCharPPI | Contact potential, calculated between intermolecular residues | Proteins 59(1):49 (2005) and BMC Bioinformatics 11:92 (2010) |

| | | | |
|---|---|---|---|
| **CP_RMFCA** | CCharPPI | The C_alpha-C_alpha potential | Proteins 65(3):726 (2006). |
| **CP_TD** | CCharPPI | Contact potential, calculated between intermolecular residues | Proteins 59(1):49 (2005) and BMC Bioinformatics 11:92 (2010) |
| **CP_SKOIP** | CCharPPI | The residue level interaction contact potential | Biophys. J. 84(3):1895 (2003). |
| **CP_D1** | CCharPPI | A reimplementation of the DECK residue level distance-dependent potential | BMC Bioinformatics. 2011 12:280. |
| **CP_Qp** | CCharPPI | Contact potential, calculated between intermolecular residues | Proteins 59(1):49 (2005) and BMC Bioinformatics 11:92 (2010) |
| **SIPPER** | CCharPPI | The SIPPER potential, described in J, Chem, Inf, Mod, 51 | J. Chem. Inf. Mod. 51:370 (2011). |
| **AP_DFIRE2** | CCharPPI | Interaction energy calculated using the DFIRE2 potential | Protein Sci. 17:1212 (2008). |
| **AP_dDFIRE** | CCharPPI | Interaction energy calculated using the dDFIRE potential | Proteins 72:793 (2008). |
| **PYDOCK_TOT** | CCharPPI | Total pyDock energy | Proteins 68:503 (2007) and Protein 69:852 (2007). |
| **CP_RMFCEN1** | CCharPPI | The 6bin-HRSC centroid-centroid potential | Proteins 70(3):950 (2006). |
| **CP_RMFCEN2** | CCharPPI | The 7bin-HRSC centroid-centroid potential | Proteins 70(3):950 (2006). |
| **CP_TSC** | CCharPPI | The residue level interaction two-step potentia | BMC Struct biol 10:40 (2010). |
| **CP_MJ2h** | CCharPPI | Contact potential, calculated between intermolecular residues | Proteins 59(1):49 (2005) and BMC Bioinformatics 11:92 (2010) |
| **CP_TB** | CCharPPI | The residue level interaction contact potential | Proteins 62(4):970 (2006). |
| **BSA_Apolar** | Freesasa | Polar Buried Surface Area from FreeSASA | F1000Research 5:189 (2016) |

| | | | |
|---|---|---|---|
| **CP_BT** | CCharPPI | Contact potential, calculated between intermolecular residues | Proteins 59(1):49 (2005) and BMC Bioinformatics 11:92 (2010) |
| **AP_DDG_W** | CCharPPI | The weighted atomic potential derived from mutation data | J. Chem. Theory Comput. 9(8):3715-3727 (2013). |
| **AP_DARS** | CCharPPI | The DARS potential | Biophys J. 2008 95(9):4217-27 |
| **FIREDOCK** | CCharPPI | The total FireDock energy (default energy function) | Proteins 69(1):139 (2007). |
| **AP_PISA** | CCharPPI | The PISA score | Proteins 81(4):592 (2013). |
| **CP_TEl** | CCharPPI | Contact potential, calculated between intermolecular residues | Proteins 59(1):49 (2005) and BMC Bioinformatics 11:92 (2010) |
| **CP_TS** | CCharPPI | Contact potential, calculated between intermolecular residues | Proteins 59(1):49 (2005) and BMC Bioinformatics 11:92 (2010) |
| **AP_DDG_U** | CCharPPI | The unweighted atomic potential derived from mutation data | J. Chem. Theory Comput. 9(8):3715-3727 (2013). |
| **CP_ZS3DC_MIN** | CCharPPI | The E_ZS3DC z-score R_min potential | Protein Sci. 2011 20(3):529-41. |
| **CP_BFKV** | CCharPPI | Contact potential, calculated between intermolecular residues | Proteins 59(1):49 (2005) and BMC Bioinformatics 11:92 (2010) |
| **PROPNSTS** | CCharPPI | Amino acid propensity score | J. Chem. Inf. Mod. 51:370 (2011). |
| **cips_AlAr** | CIPS | Sum of CIPS score for Aliphatic-Aromatic contacts | Bioinformatics 34:459 (2018) |
| **AlAr** | COCOMAPS | Aliphatic-Aromatic contact count at 5 Å distance | Bioinformatics 27:2915 (2011) |
| **CP_MJPL** | CCharPPI | Contact potential, calculated between intermolecular residues | Proteins 59(1):49 (2005) and BMC Bioinformatics 11:92 (2010) |

| | | | |
|---|---|---|---|
| **CP_SKOb** | CCharPPI | Contact potential, calculated between intermolecular residues | Proteins 59(1):49 (2005) and BMC Bioinformatics 11:92 (2010) |
| **AP_MPS** | CCharPPI | The MPS potential | Biophys J. 2008 95(9):4217-27 |
| **ArAr** | COCOMAPS | Aromatic-Aromatic contact count at 5 Å distance | Bioinformatics 27:2915 (2011) |
| **RFC** | This work | Random forest classifier | |
| **NNC** | This work | Perceptron classifer | |
| **TF** | This work | Deep neural network classifer | |

**Table S3.** Summary report from snorkel summary report for the labelling functions. We generated labeling functions using the top 30 scoring functions which had varying accuracies ranging from 66% to 84% (Initial optimization). For each scoring function, we aimed to find the thresholds that maximize the true positive rate (TPR) and the true negative rate (TNR). Optimizing snorkel using this approach resulted in modest results, achieving about 74% labeling accuracy on the training dataset and about 70% labeling accuracy on the validation dataset. The results had a small variance when training snorkel with either top 10, top 20, or top 30 scoring functions. The final set of labelling functions (Final optimization) presented better accuracy on the train and testing set (86.26% and 87.71% respectively ) at the cost of reducing the coverage but avoiding the high overlap presented on the initial optimization.

| Initial optimization | j | Polarity | Coverage | Overlaps | Conflicts | Correct | Incorrect | Emp. Acc. |
|---|---|---|---|---|---|---|---|---|
| lf_CONSRANK_val | 0 | [0, 1] | 0.999982 | 0.999982 | 0.924374 | 47025 | 8722 | 0.843543 |
| lf_CP_HLPL | 1 | [0, 1] | 0.995516 | 0.995516 | 0.919907 | 39622 | 15876 | 0.713936 |
| lf_CP_MJ3h | 2 | [0, 1] | 0.998726 | 0.998726 | 0.923118 | 39677 | 16000 | 0.712628 |
| lf_DDG_V | 3 | [0, 1] | 0.999982 | 0.999982 | 0.924374 | 39669 | 16078 | 0.711590 |
| lf_CP_RMFCA | 4 | [0, 1] | 0.999982 | 0.999982 | 0.924374 | 39463 | 16284 | 0.707895 |
| lf_AP_GOAP_DF | 5 | [0, 1] | 0.999982 | 0.999982 | 0.924374 | 39757 | 15990 | 0.713168 |
| lf_CP_Qp | 6 | [0, 1] | 0.993973 | 0.993973 | 0.918437 | 38619 | 16793 | 0.696943 |
| lf_CP_TD | 7 | [0, 1] | 0.999193 | 0.999193 | 0.923603 | 39688 | 16015 | 0.712493 |
| lf_CP_SKOIP | 8 | [0, 1] | 0.998099 | 0.998099 | 0.922490 | 39032 | 16610 | 0.701484 |
| lf_CP_TB | 9 | [0, 1] | 0.999856 | 0.999856 | 0.924248 | 38197 | 17543 | 0.685271 |
| lf_CP_TSC | 10 | [0, 1] | 0.999677 | 0.999677 | 0.924087 | 38290 | 17440 | 0.687063 |
| lf_PYDOCK_TOT | 11 | [0, 1] | 0.999982 | 0.999982 | 0.924374 | 38090 | 17657 | 0.683265 |
| lf_SIPPER | 12 | [0, 1] | 0.999928 | 0.999928 | 0.924320 | 38705 | 17039 | 0.694335 |

| | | | | | | | | |
|---|---|---|---|---|---|---|---|---|
| lf_CP_BT | 13 | [0, 1] | 0.998404 | 0.998404 | 0.922795 | 37567 | 18092 | 0.674949 |
| lf_CP_MJ2h | 14 | [0, 1] | 0.999856 | 0.999856 | 0.924248 | 38440 | 17300 | 0.689630 |
| lf_AP_DFIRE2 | 15 | [0, 1] | 0.999982 | 0.999982 | 0.924374 | 38833 | 16914 | 0.696594 |
| lf_CP_RMFCEN1 | 16 | [0, 1] | 0.999767 | 0.999767 | 0.924159 | 38617 | 17118 | 0.692868 |
| lf_AP_DARS | 17 | [0, 1] | 1.000000 | 1.000000 | 0.924392 | 37980 | 17768 | 0.681280 |
| lf_AP_PISA | 18 | [0, 1] | 1.000000 | 1.000000 | 0.924392 | 37548 | 18200 | 0.673531 |
| lf_BSA_Apolar | 19 | [0, 1] | 1.000000 | 1.000000 | 0.924392 | 38100 | 17648 | 0.683433 |
| lf_CP_BFKV | 20 | [0, 1] | 0.999964 | 0.999964 | 0.924356 | 37245 | 18501 | 0.668120 |
| lf_AP_dDFIRE | 21 | [0, 1] | 0.999265 | 0.999265 | 0.923674 | 38233 | 17474 | 0.686323 |
| lf_CP_RMFCEN2 | 22 | [0, 1] | 0.999785 | 0.999785 | 0.924177 | 38274 | 17462 | 0.686702 |
| lf_CP_ZS3DC_MIN | 23 | [0, 1] | 1.000000 | 1.000000 | 0.924392 | 36264 | 19484 | 0.650499 |
| lf_AP_DDG_U | 24 | [0, 1] | 0.999982 | 0.999982 | 0.924374 | 36939 | 18808 | 0.662619 |
| lf_AP_DDG_W | 25 | [0, 1] | 0.999982 | 0.999982 | 0.924392 | 37698 | 18049 | 0.676234 |
| lf_cips_AlAr | 26 | [0, 1] | 1.000000 | 1.000000 | 0.924392 | 36932 | 18816 | 0.662481 |
| lf_CP_MJPL | 27 | [0, 1] | 0.998888 | 0.998888 | 0.923334 | 37111 | 18575 | 0.666433 |
| lf_CP_SKOb | 28 | [0, 1] | 0.987856 | 0.987856 | 0.912553 | 36336 | 18735 | 0.659803 |
| lf_CP_TEl | 29 | [0, 1] | 0.999354 | 0.999354 | 0.923764 | 37411 | 18301 | 0.671507 |

| Final optimization | j | Polarity | Coverage | Overlaps | Conflicts | Correct | Incorrect | Emp. Acc. |
|---|---|---|---|---|---|---|---|---|
| lf_CONSRANK_val | 0 | [0, 1] | 0.218232 | 0.174141 | 0.010494 | 11898 | 268 | 0.977971 |
| lf_CP_HLPL | 1 | [0, 1] | 0.244583 | 0.210106 | 0.005058 | 11659 | 1976 | 0.855079 |
| lf_CP_MJ3h | 2 | [0, 1] | 0.231937 | 0.189011 | 0.008825 | 11166 | 1764 | 0.863573 |
| lf_DDG_V | 3 | [0, 1] | 0.238197 | 0.210716 | 0.007857 | 11351 | 1928 | 0.854808 |
| lf_CP_RMFCA | 4 | [0, 1] | 0.236690 | 0.211685 | 0.006224 | 11192 | 2003 | 0.848200 |
| lf_AP_GOAP_DF | 5 | [0, 1] | 0.226645 | 0.198805 | 0.009041 | 11157 | 1478 | 0.883023 |
| lf_NNC | 6 | [0, 1] | 0.211254 | 0.197047 | 0.004377 | 11723 | 54 | 0.995415 |
| lf_TF | 7 | [0, 1] | 0.215165 | 0.210788 | 0.003946 | 11914 | 81 | 0.993247 |

**Supplementary Section 4**. Classifiers details and hyperparameter

Perceptron

The hyperparameters for the perceptron consisted on the learning rate (alpha) of 3.7e-06 ,a batch size of 512 samples , and one hidden layer of 24 neurons.

Random Forest Classifier

This classifiers parameters consisting on 100 classifiers to make the forest, a minimum number of samples to split the node of ten, and weighted classes (Incorrect: 1.2 and correct : 0.1 ).

Deep neural network

We trained a sequential deep neural network using Tensorflow 2.0 framework and keras. We combined dense and dropout layers, the latter to drop 20% of the neurons. This model has a total of 2,665 trainable parameters.

| Type | Output Shape | Param num |
|---|---|---|
| Dense | (None, 40) | 360 |
| Dense | (None, 16) | 656 |
| Dropout | (None, 16) | 0 |
| Dense | (None, 16) | 272 |
| Dense | (None, 16) | 272 |
| Dropout | (None, 16) | 0 |
| Dense | (None, 16) | 272 |
| Dense | (None, 16) | 272 |
| Dropout | (None, 16) | 0 |
| Dense | (None, 16) | 272 |
| Dense | (None, 16) | 272 |
| Dense | (None, 1) | 17 |

**Table S5.** Success rate on the **3K-BM5up** for all the 20 best performing classifiers/scoring functions in terms of percentage of targets featuring at least one correct solution within the top-10 ranked positions. Values for the top-1 and top-100 positions are also reported.

| | **3K-BM5up** | | |
|---|---|---|---|
| **Method** | **Top-1** | **Top-10** | **Top-100** |
| **RF-hAIkal** | 50.0 | 55.8 | 69.2 |
| **PRC-hAIkal** | 43.6 | 54.6 | 69.1 |
| **RF-core** | 32.7 | 54.6 | 69.1 |
| **PRC-core** | 43.6 | 50.9 | 56.4 |
| CP_TSC | 14.6 | 49.1 | 65.4 |
| **TF-core** | 30.9 | 43.6 | 65.4 |
| **TF-hAIkal** | 29.1 | 41.8 | 69.1 |
| AP_PISA | 21.8 | 41.8 | 67.3 |
| CP_BT | 20.0 | 40.0 | 60.0 |
| CP_TB | 16.4 | 40.0 | 63.6 |

| | | | |
|---|---|---|---|
| CP_MJ2h | 14.6 | 40.0 | 63.6 |
| CP_Qp | 12.7 | 40.0 | 67.3 |
| CP_TS | 9.1 | 40.0 | 67.3 |
| CP_HLPL | 18.2 | 38.2 | 69.1 |
| CP_BFKV | 18.2 | 36.4 | 61.8 |
| CP_TD | 9.1 | 36.4 | 54.6 |
| CONSRANK_val | 32.7 | 34.6 | 41.8 |
| CP_MJ3h | 12.7 | 34.6 | 58.2 |
| DDG_V | 16.4 | 32.7 | 60.0 |

**Table S6.** Success rate on the **Score_set** for all the 20 best performing classifiers/scoring functions in terms of percentage of targets featuring at least one correct solution within the top-10 ranked positions. Values for the top-1 and top-100 positions are also reported.

| **Score_set** | | | |
|---|---|---|---|
| **Method** | **Top-1** | **Top-10** | **Top-100** |
| **TF-hAIkal** | 26.7 | 66.7 | 66.7 |
| AP_PISA | 40.0 | 53.3 | 66.7 |
| **RF-hAIkal** | 33.3 | 53.3 | 60.0 |
| **TF-core** | 20.0 | 53.3 | 66.7 |
| **PRC-core** | 40.0 | 46.7 | 53.3 |
| **RF-core** | 33.3 | 46.7 | 53.3 |
| CP_TB | 26.7 | 46.7 | 80.0 |
| CP_BT | 13.3 | 46.7 | 60.0 |
| CP_TSC | 13.3 | 46.7 | 73.3 |
| CP_D1 | 13.3 | 46.7 | 66.7 |
| **PRC-hAIkal** | 26.7 | 40.0 | 53.3 |
| CONSRANK_val | 26.7 | 40.0 | 46.7 |
| SIPPER | 6.7 | 40.0 | 66.7 |
| CP_BFKV | 6.7 | 40.0 | 66.7 |
| CP_TEl | 6.7 | 40.0 | 66.7 |

| | | | |
|---|---|---|---|
| DDG_V | 13.3 | 33.3 | 53.3 |
| PYDOCK_TOT | 13.3 | 33.3 | 46.7 |
| AP_dDFIRE | 6.7 | 33.3 | 60.0 |
| PROPNSTS | 6.7 | 33.3 | 73.3 |
| AP_DFIRE2 | 13.3 | 26.7 | 66.7 |

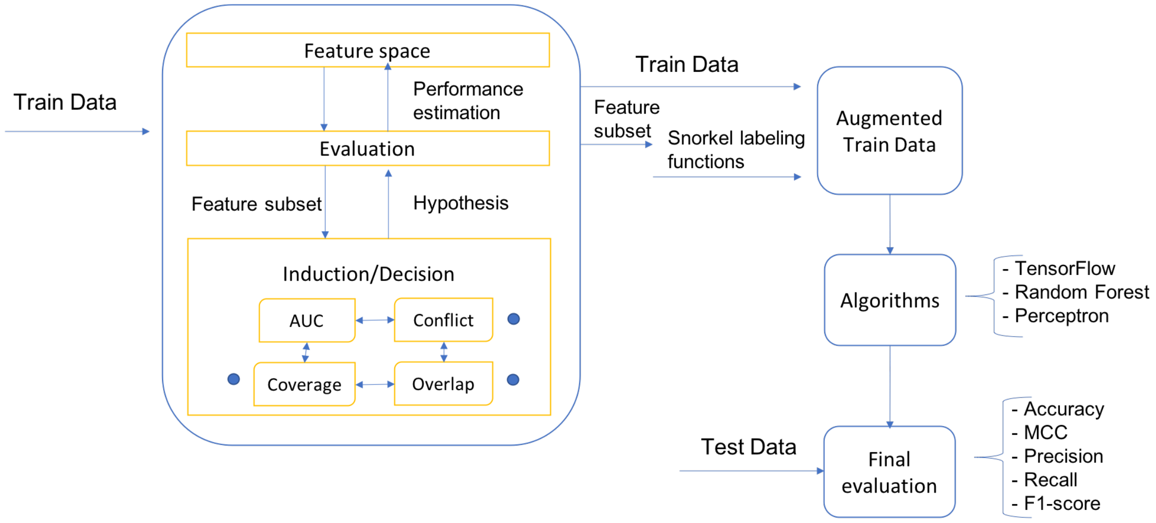

**Fig. S6:** Feature selection process. From out train date , we wrap the feature selection process, in different steps. starting from all the features available, we evaluated this features using the AUC value and the snorkel built-in evaluation metrics overlap and conflict. We selected the best features to train different algorithms based on high AUC, low conflict , low overlap and certain coverage. The next step was to obtain the final evaluation with our selected features and metrics.

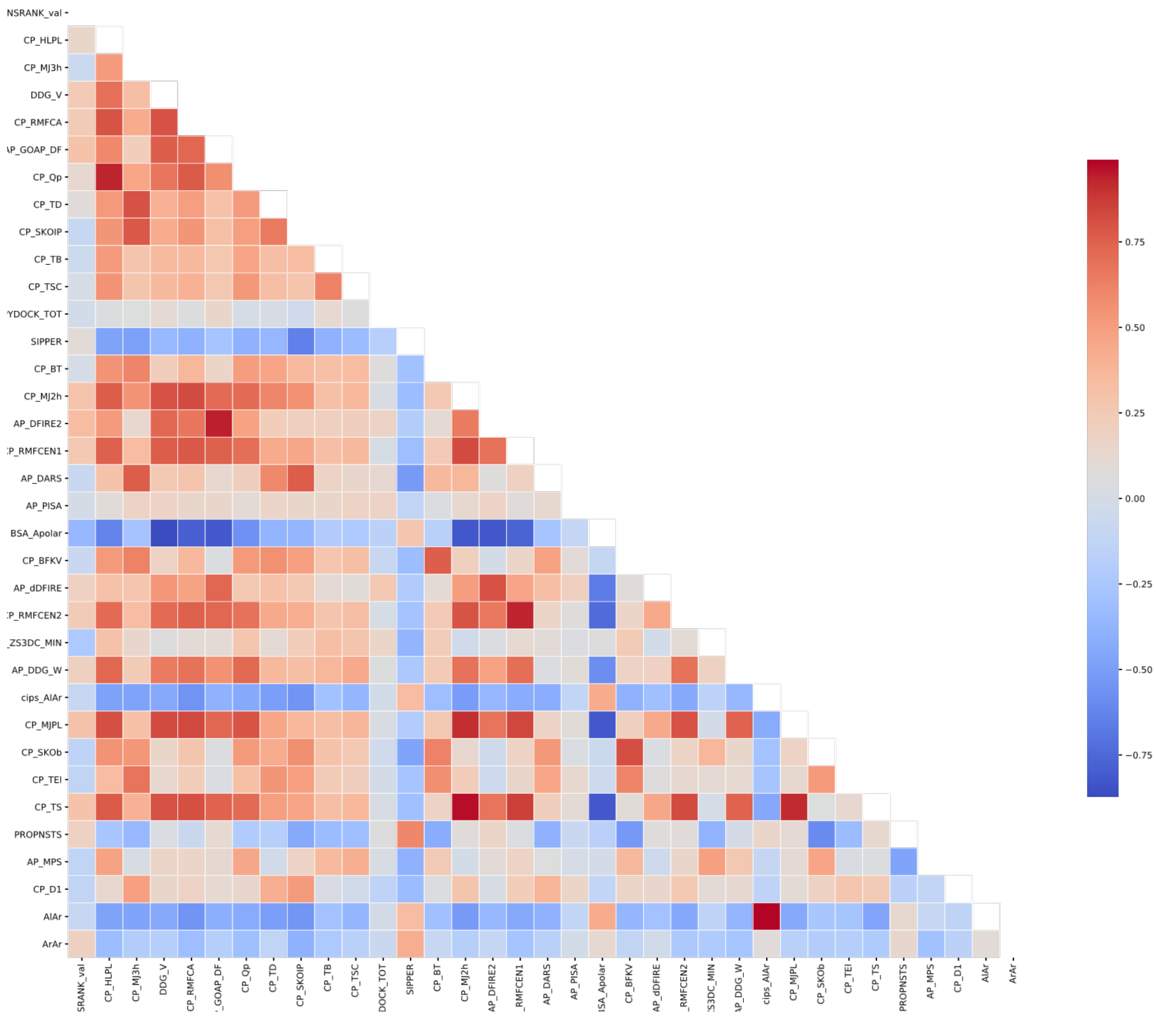

**Fig. S7:** Pearson correlation of top performing scoring functions according to the ROC on the balanced training data. The linear correlation of the scoring function is indicative of the usefulness to separate the data. The values are in blue-white read color scale, where the lowest correlation have the colorblue, and the highest color red
